# A Deficiency in SUMOylation Activity Disrupts Multiple Pathways Leading to Neural Tube and Heart Defects in *Xenopus* Embryos

**DOI:** 10.1101/549105

**Authors:** Michelle M. Bertke, Kyle M. Dubiak, Laura Cronin, Erliang Zeng, Paul W. Huber

## Abstract

**Background:** Adenovirus protein, Gam1, triggers the proteolytic destruction of the E1 SUMO-activating enzyme. Microinjection of an empirically determined amount of Gam1 mRNA into one-cell *Xenopus* embryos can reduce SUMOylation activity to undetectable, but nonlethal, levels, enabling an examination of the role of this post-translational modification during early vertebrate development.

**Results:** We find that SUMOylation-deficient embryos consistently exhibit defects in neural tube and heart development. We have measured differences in gene expression between control and embryos injected with Gam1 mRNA at three developmental stages: early gastrula (immediately following the initiation of zygotic transcription), late gastrula (completion of the formation of the three primary germ layers), and early neurula (appearance of the neural plate). Although changes in gene expression are widespread and can be linked to many biological processes, three pathways, non-canonical Wnt/PCP, snail/twist, and Ets-1, are especially sensitive to the loss of SUMOylation activity and can largely account for the predominant phenotypes of Gam1 embryos. SUMOylation appears to generate different pools of a given transcription factor having different specificities with this post-translational modification involved in the regulation of more complex, as opposed to housekeeping, processes.

**Conclusions:** We have identified changes in gene expression that underlie the neural tube and heart phenotypes resulting from depressed SUMOylation activity. Notably, these developmental defects correspond to the two most frequently occurring congenital birth defects in humans, strongly suggesting that perturbation of SUMOylation, either globally or of a specific protein, may frequently be the origin of these pathologies.

## Background

Post-translational modification by small ubiquitin-like modifier (SUMO) has emerged as a global mechanism for the regulation of protein activity, stability, and localization [1–5]. Its integration into other types of protein modification, such as phosphorylation, acetylation, and ubiquination, allows for coordinated control of diverse biological processes. Although the SUMO (E2) conjugating enzyme is capable of acting directly on target proteins, in the majority of cases a sizable family of SUMO (E3) ligases determine substrate specificity. The SENP family of proteases, likewise exhibiting different specificities, acts to remove SUMO moieties so that the modification is reversible and can be tightly regulated.

The steady-state level of SUMOylated protein in any given instance is typically low and has been referred to as the “SUMO enigma” [6]. This has made detection of SUMO-modified proteins difficult. Nonetheless, a strategy that employed the expression of epitope-tagged SUMO2 in human cells combined with flow cytometry enabled the identification of more than 1,600 proteins conjugated to this SUMO isoform [4]. Similar to earlier studies with yeast cells [5, 7], SUMO modification in this case was mostly limited to nuclear proteins or those that are nucleocytoplasmic. This is consistent with the large number of transcription factors, RNA binding proteins, chromatin associated proteins, and cell cycle regulators that are documented targets of SUMOylation [8, 9]. In *Xenopus* egg extract, a large number of SUMOylated proteins are associated with chromatin [10]. However, a bioinformatic analysis of this proteomic data revealed that an appreciable number (39.5%) of the identified SUMOylated proteins were linked to metabolic processes and translation, indicating the importance of this post-translational modification in cytoplasmic processes as well.

The importance and scope of SUMOylation during early development has been difficult to ascertain, since complete elimination of this activity is embryonic lethal [2, 8, 11–13]. From yeast to vertebrates, the essential role of SUMOylation in mitotic processes has defeated approaches that have knocked out this pathway by elimination of the sole E2 conjugating enzyme. Strategies that have eliminated only one of the SUMO isoforms have resulted in less severe phenotypes in the case of SUMO1 [14–16], but embryonic lethal phenotypes in the case of SUMO2 [17]. Gene knockout or other methods of gene inactivation that target other components of the SUMOylation machinery (*i.e.,* E3 ligases and SENP proteases) result in a variety of phenotypes [11].

A means to diminish, but not eliminate, SUMOylation activity would enable an assessment of the pathway’s importance and ubiquity during early development. The avian adenovirus protein, Gam1, binds to the SAE1 subunit of the E1 SUMO-activating enzyme and triggers its proteolytic degradation [18, 19], and injection of mRNA encoding Gam1 into *Xenopus* embryos reduces SUMOylation activity to undetectable levels [20]. Here, we have used Gam1 in order to measure changes in gene expression in SUMOylation-deficient embryos at three developmental stages, early gastrula, late gastrula, and early neurula using microarray technology, which can analyze the activity of approximately 30,000 genes. The data show that SUMOylation impacts many disparate biological processes; nonetheless, certain signaling pathways appear to be particularly sensitive to depletion of this activity and can be correlated with the predominant phenotypes of these embryos that include failure of blastopore and neural tube closure, shortened anterior-posterior (A-P) axis, and defective heart and eye development. Importantly, we show that disruptions in these pathways due to loss of this post-translational modification during the earliest periods of embryogenesis can elicit developmental deficits that correspond to the most frequently occurring human birth defects.

## Results and Discussion

### Gam1 depletion of SUMOylation activity in Xenopus embryos

The adenovirus protein Gam1 triggers the proteolytic destruction of the SUMO E1 activating enzyme [18, 19], providing a strategic alternative to antisense methods, since the effect is immediate and eliminates both existing and *de novo* accumulation of the activating enzyme. We have reported that injection of mRNA encoding Gam1 into one-cell *Xenopus* embryos suppresses SUMOylation activity to undetectable levels in extract prepared from early neurula stage embryos [20]. In order to characterize the activity of Gam1 further, we measured the persistence of the protein during development. Fertilized eggs were injected with mRNA (0.5 ng) encoding myc-tagged Gam1; protein extract prepared from embryos at the designated developmental stage was analyzed by western blot. Gam1 protein is present by midblastula stage and reaches a maximum during early gastrula before declining during neurulation (Fig. 1A).

**Figure 1.**
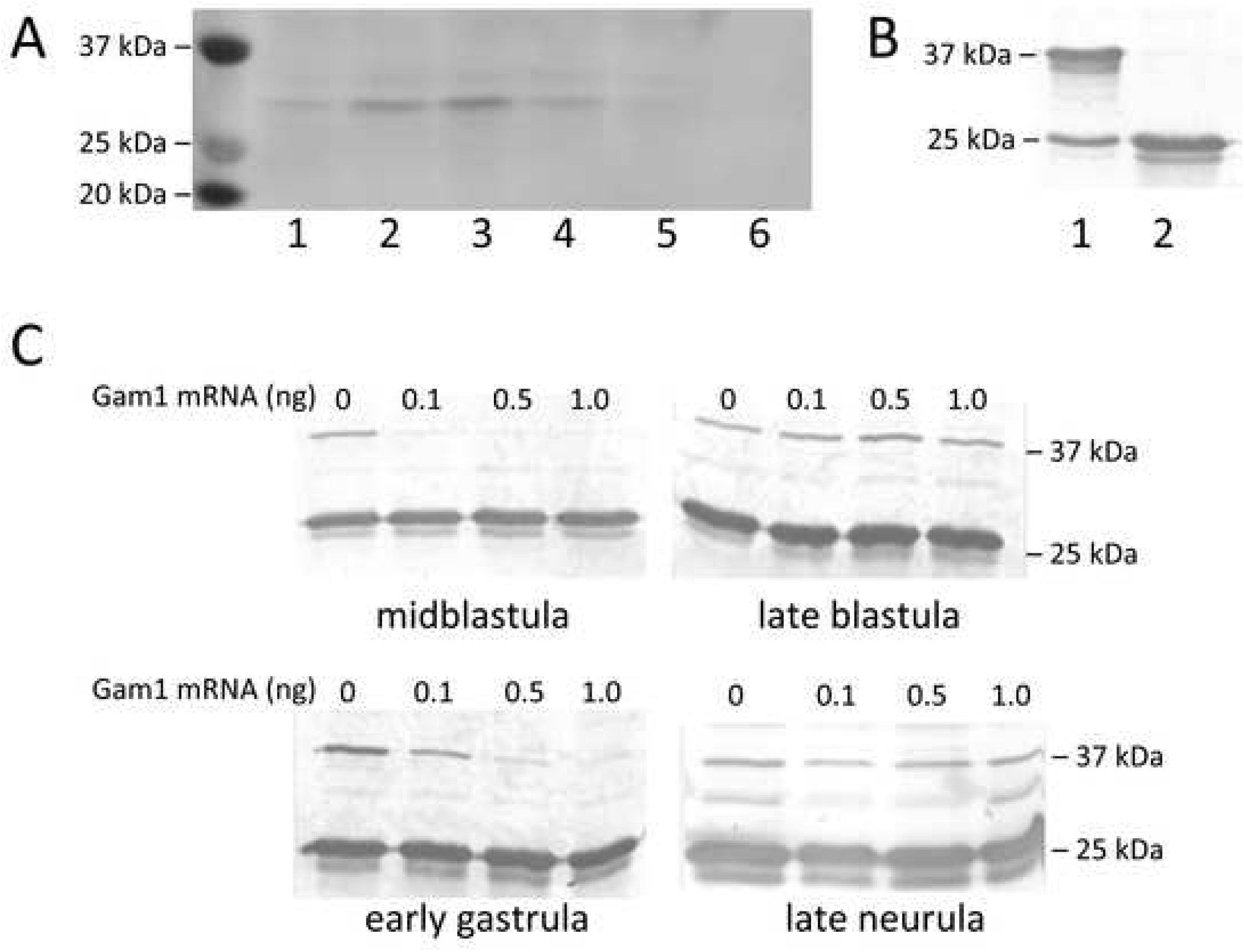
Depletion of SUMOylation activity in embryos injected with Gam1 mRNA. (A) Expression levels of Gam1 protein during embryogenesis. One-cell embryos were injected with mRNA (0.5 ng) encoding Gam1 with an N-terminal myc tag. Whole cell protein extract was prepared from embryos at the indicated Nieuwkoop-Faber stage and 25 μg taken for western blot analysis using anti-myc antibody. *Lanes:* 1, midblastula; 2, late blastula; 3, early gastrula; 4, early neurula; 5, late neurula; 6, water injected control. (B) *In vitro* SUMOylation assay. All assays contained Ubc9 (E2 enzyme), SUMO1, ATP, and substrate peptide (25 kDa) with (lane 1) or without (lane 2) purified E1 enzyme (500 nM) added. Samples were analyzed by western blot using an antibody specific for a 25 kDa SUMO substrate peptide. (C) Assays for E1 activity. One-cell embryos were injected with the indicated amount of Gam1 mRNA and allowed to develop to the indicated stage. Whole cell extract was prepared from 20 embryos and an equivalent amount of protein (25 ug) was used as a source of E1 enzyme for each assay.

An *in vitro* assay containing E1, E2 (UBC9), SUMO1, and a 25 kDa substrate peptide was used to measure SUMOylation activity by western blot (Fig. 1B). The effect of different amounts of injected Gam1 mRNA on SUMOylation activity was determined at four stages of development using whole cell extract prepared from water or Gam1 injected embryos as a source of E1 enzyme in the *in vitro* assay (Fig. 1C). All three amounts of injected Gam1 mRNA effectively suppress SUMOylation activity at midblastula. However, at late blastula there is measureable SUMOylation activity in all samples, presumably due to the approximate 2-fold increase during mid- and late blastula in the mRNAs that encode the two subunits (SAE and UBA2) that comprise the E1 enzyme [21]. Suppression of SUMOylation activity is reestablished in early gastrula embryos (0.5 or 1 ng injected mRNA) that then persist until late neurula. The excellent correlation between the levels of Gam1 protein and suppression of E1 activity is in accord with other evidence that expression of the viral protein is an effective method to control the SUMO pathway [18, 19]. The transient appearance of SUMOylation activity at the midblastula transition (MBT) possibly accounts for the survival of these embryos. Injection of higher amounts (5 ng) of Gam1 mRNA is lethal with no survival of injected embryos beyond the gastrula-neurula transition (stage 13). A mutant form of Gam1 that cannot bind to the SAE1 subunit had no effect on viability when injected at the same amount. Based on these results, 0.5 ng of injected Gam1 mRNA was chosen for the subsequent experiments to examine the role of SUMOylation in developing embryos.

### Distinct phenotypes in SUMOylation deficient Xenopus embryos

Embryos injected with Gam1 mRNA develop normally up through late blastula compared to water injected or uninjected control embryos. One of the earliest phenotypes of Gam1 embryos is delayed or incomplete closure of the blastopore, which is a hallmark of gastrulation (Fig. 2A). In instances where closure is simply slowed, the embryo usually develops normally after this point. Embryos, in which the blastopore fails to close, do not develop normally and exhibit cascading effects that are apparent at the neurula stage of development. During neurulation, the neural tube forms due to convergent extension of the outermost layer of cells on the embryo [22, 23]. The posterior portion of the neural tube closes at or very near to the point of blastopore closure. Therefore, when the blastopore fails to close properly, the neural tube also fails to close properly and embryos display a *spina bifida* like phenotype at the late neurula stage. In the most extreme cases, embryos injected with Gam1 present completely open neural tubes with failure of the neural folds to fuse together along the entire A-P axis (Fig. 2B).

**Figure 2.**
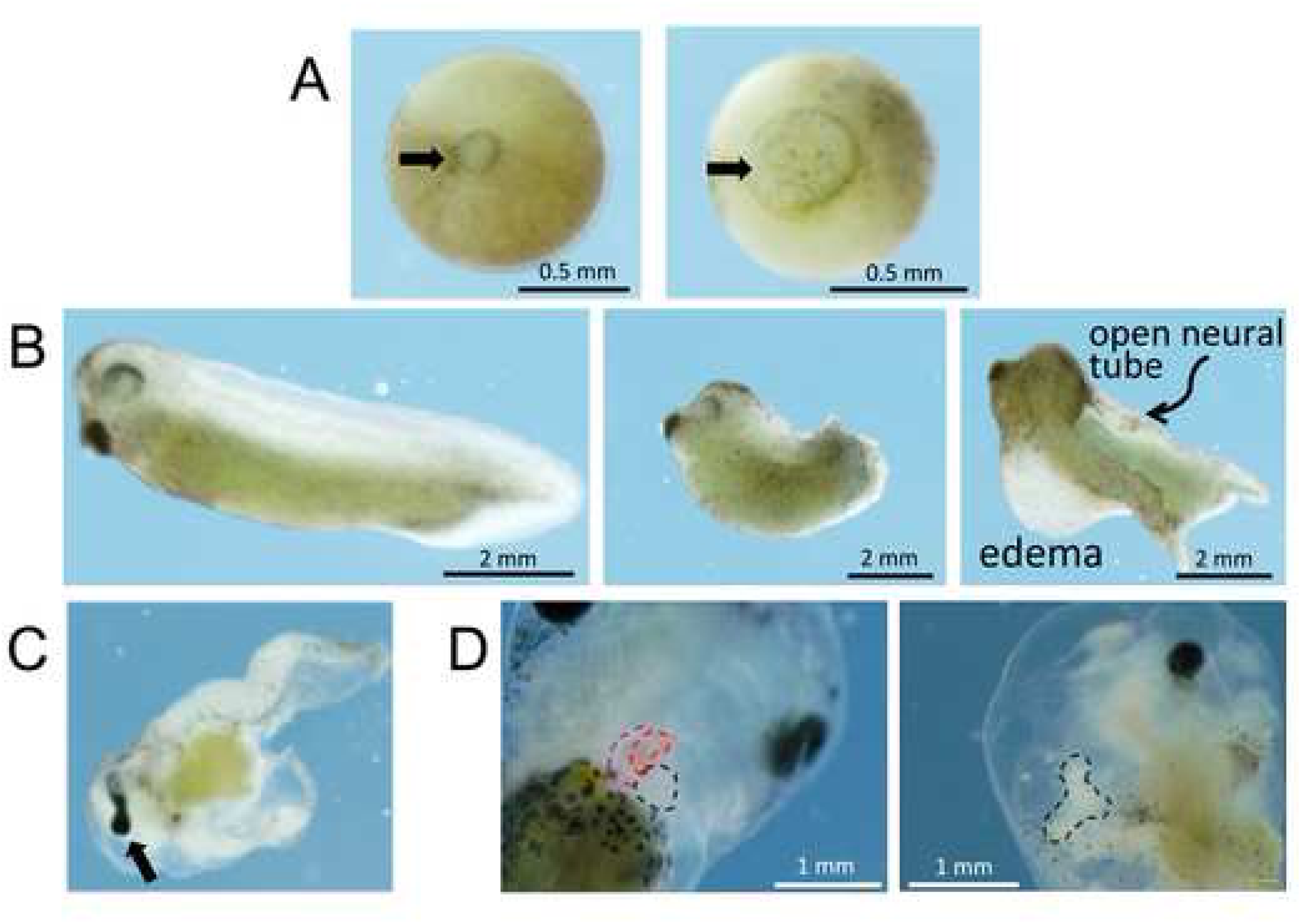
Developmental defects of Gam1-injected embryos. (A) Control *(left*) and Gam1-injected (*right*) late gastrula stage embryos showing delayed closure of the blastopore (arrow) in the latter. (B) Stage 35 (50 hpf) control embryo (*left*), Gam1 embryo with shortened A-P axis (*center*), Gam1 embryo with an open neural tube and edema around the heart (*right*). (C) An example of an eye defect (cyclopia). (D) Failure of the heart tube to initiate looping. (*Left*) A control embryo (96 hpf) in which the two atria (red) and ventricle (black) have fully developed compared to (*right*) a Gam1 embryo in which the heart tube has failed to undergo looping.

Embryos that are viable at the late neurula stage usually develop into free-swimming tadpoles (>48 hpf). Gam1 injected embryos at this stage of development have shortened or bent axes often due to the failure of the neural tube to close properly. Additionally, the majority of embryos display some degree of edema, or fluid collection, in the ventral regions of the embryo specifically around the heart and digestive system (Fig. 2B). Along with edema and a shortened axis, embryos display defects in eye, heart, and gut development, including cyclopia or eyes that are not fully separate (fused eye) (Fig. 2C) and disordered digestive system.

Defects in the heart include improper looping, disruption of chamber formation, and an absence of blood flow. Normal amphibian hearts contain three chambers, two atria and one ventricle, with looping of the outflow track to the right. Hearts of Gam1-injected embryos often lack discernible chambers and seem to have not developed beyond a heart tube structure due to a failure of proper looping (Fig. 2D). Heart structures that have formed contain little to no blood past the stage in development when control matched embryos have begun producing blood. Surprisingly, these embryos are viable and have heart structures that continue to beat, although contractions are erratic. Defects in the formation of these organs indicate that the temporary disruption of SUMOylation activity during early embryogenesis impacts the developmental program long after this activity is restored.

### Inhibition of SUMOylation activity disrupts convergence and extension

The phenotypes of a shortened A-P axis and failure of the blastopore and neural tube to close in the Gam1 embryos indicate that SUMOylation activity is necessary for convergence and extension. Information concerning the involvement of SUMO in cell movement is limited. A morpholino oligonucleotide directed at SUMO1 prevented activin-induced elongation of *Xenopus* animal caps [15] and SUMOylation of Rac1 GTPase is needed for optimal cell migration in response to hepatocyte growth factor [24]. Since cell movement during gastrulation is primarily internal and cannot be easily observed, we turned to Keller explant assays to determine whether the SUMOylation deficiency in Gam1 embryos impacts this activity [25]. Explants are made by sandwiching together a portion of two embryos dorsal to the blastopore lip, which contain migrating cells. This arrangement causes the mesodermal and endodermal cells to elongate in a plane rather than involuting beneath ectodermal cells, enabling quantification of the degree of convergent extension.

Control embryos (injected with H_2_O) showed normal convergent extension while those injected with 2.5 ng Gam1 mRNA failed to elongate properly (Fig. 3A). Additionally, when the amount of Gam1 mRNA was doubled to 5.0 ng, explants not only showed no evidence of cell movement, but the two explants failed to adhere to each other. This indicates not only a role of SUMOylation in controlling cell migration, but also in controlling cell adhesion.

**Figure 3.**
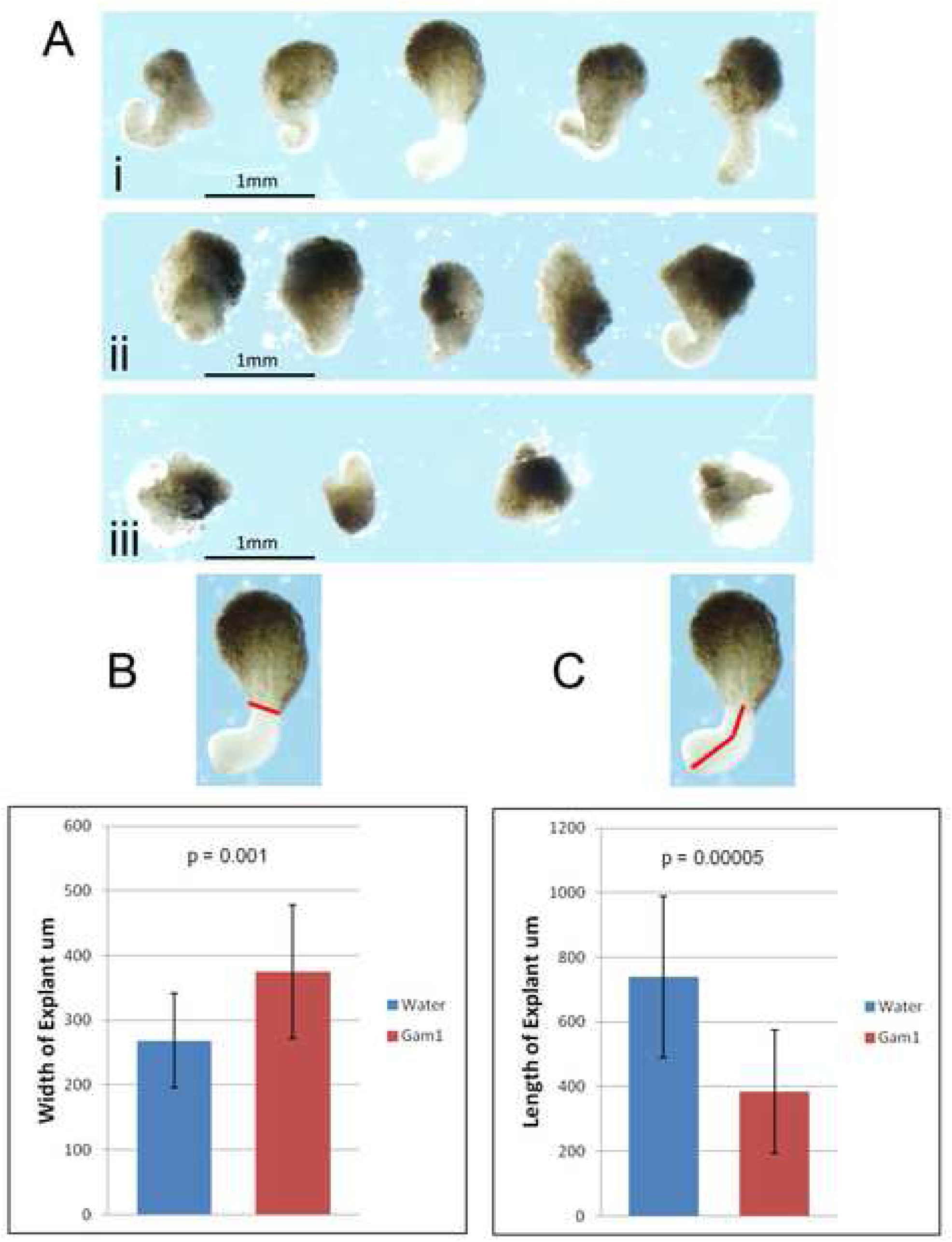
Gam1 disrupts convergence and extension. (A) Examples of Keller sandwich explants prepared from embryos injected with (i) water, (ii) 2.5 ng Gam1 mRNA, (iii) 5 ng Gam1 mRNA. Explants from embryos injected with water (n = 17) or 2.5 ng Gam1 mRNA (n = 18) were measured to quantify effects on (B) convergence and (C) extension. The *red* lines designate the lengths measured in the explants. Error bars represent standard deviation.

The degree of explant convergent extension was quantified by measuring the boundary of the non-involuting and involuting marginal zones (convergence) and the anterior to posterior length (extension) (Fig. 3B,C). Two-tailed t-tests show that both convergence (p < 0.001) and extension (p < 0.00005) are significantly decreased in Gam1 embryos compared to controls. The disruption of these activities explains, at least in part, many of the observed phenotypes of Gam1 embryos.

### Gene expression in SUMOylation deficient Xenopus embryos

Gam1 knockdown of SUMOylation activity reproducibly generated distinct developmental phenotypes, several of which have been previously linked to disruptions of this post-translational modification, including neural tube closure [26], cleft lip/palate [27], heart defects [28–30], axial mesodermal defects that lead to midline phenotypes such as cyclopia [31], and disruption of hematopoiesis [32]. In good accord with the Gam1 phenotypes, a morpholino antisense oligonucleotide directed against *Xenopus* SUMO1 caused shortened and bent A-P axes, incomplete closure of the neural tube, microcephaly, and inhibition of activin-induced elongation of animal caps [15]. The close correspondence between the morphlino and Gam1 phenotypes affirm that the effects resulting from Gam1 expression are chiefly due to interference of SUMOylation activity in these embryos.

The ability to decrease SUMOylation activity to nonlethal levels presents the opportunity to examine the role of this protein modification, which frequently targets transcription factors and several other nuclear proteins, in the regulation of gene expression during early development. Three developmental stages were chosen for transcriptome analysis (30,000 genes) by microarray: early gastrula (stage 10), which shortly follows the initiation of zygotic transcription at the MBT; late gastrula (stage 12), which is at the end of a period of complex cell movement that organizes the three germ layers; and early neurula (stage 14), which marks changes in transcription that underlie organogenesis and patterning of the embryo. Three biological replicates were analyzed in order to account for variation between egg clutches. The R-values of scatter plots of the biological replicates ranged from 0.959 to 0.916, demonstrating excellent correlation between each of the replicates. The number of differentially expressed genes (p < 0.05) between control (water injected) and Gam1 embryos numbers 94 (53 down-regulated, 41 up-regulated) at early gastrula; 447 (263 down-regulated, 221 up-regulated) at late gastrula; and 742 (394 down-regulated, 387 up-regulated) at early neurula. A heatmap of all differentially expressed genes identified from comparisons between control and Gam1 injected embryos at the three developmental stages is presented in Additional file 1: Fig. S1.

In order to validate the microarray data, the same RNA samples were used for qRT-PCR assays of genes selected based upon either the magnitude of the change in expression (*dmrta1*, *rpl8*, *ccng1*), the occurrence of the gene in a pathway involved in patterning the early embryo (*xbra*, *foxc1*, *wnt8b*, *gsc*, *chrd*), or a previous microarray analysis of *X. laevis* development (*xpo1*, *wnt8b*, *krt*) [33]. (A graphical comparison of the microarray and qRT-PCR data is presented in Additional file 2: Fig. S2). The expected correlation between microarray and qRT-PCR is 80 - 87% with respect to the direction of the change (up-*versus* down-regulation) [33, 34]. The correlation between our microarray and qRT-PCR assays is 93%; these results, along with the scatter plots, indicate that the quality of the microarray data is high. There is also good agreement between the two measurements with regard to quantitation, especially for those genes that show the greatest up-(*ccng1* and *xpo*) and down-regulation (*dmrta1*).

### Biological process ontology of differentially expressed genes

Differentially regulated genes for each time point were assembled into gene lists for analysis using data mining software including MetaCore, BiNGO, DAVID, and the Gene Ontology database [35]. Volcano plots (Additional file 3: Fig. S3) display statistical significance (*p* value) versus fold change at each time point. These plots reflect not only the increasing number of differentially expressed genes progressing from early gastrula to early neurula, but also the increasing magnitude of these differences. With few exceptions, most changes in expression at early gastrula are only 20 - 30%; whereas, larger quantitative changes are seen at the later two time points. We have used these lists of genes affected by the loss of SUMOylation activity for various bioinformatics analyses in order to understand the regulatory role of this post-translational modification during early vertebrate development.

The first analysis was to determine the predominant (high level) biological processes affected by loss of SUMOylation activity at the three different developmental time points (Fig. 4). The profiles at these three stages, which span a total of about 7 hours, are distinctly different. However, the distribution of affected genes among these biological processes seems to reflect the major cellular activities of embryos at each of these stages. With lengthening of the cell cycle, zygotic transcription begins at the MBT and transcriptome analysis at early gastrula is expected to capture this transition. Indeed, many of the differentially expressed genes at this time point normally show robust induction between Nieuwkoop-Faber stages 8 and 10 [36]. The greatest proportion of affected genes at early gastrula are concerned with Regulation (31%) and Metabolism (18%), primarily as a result of the resumption of transcription and protein synthesis.

**Figure 4.**
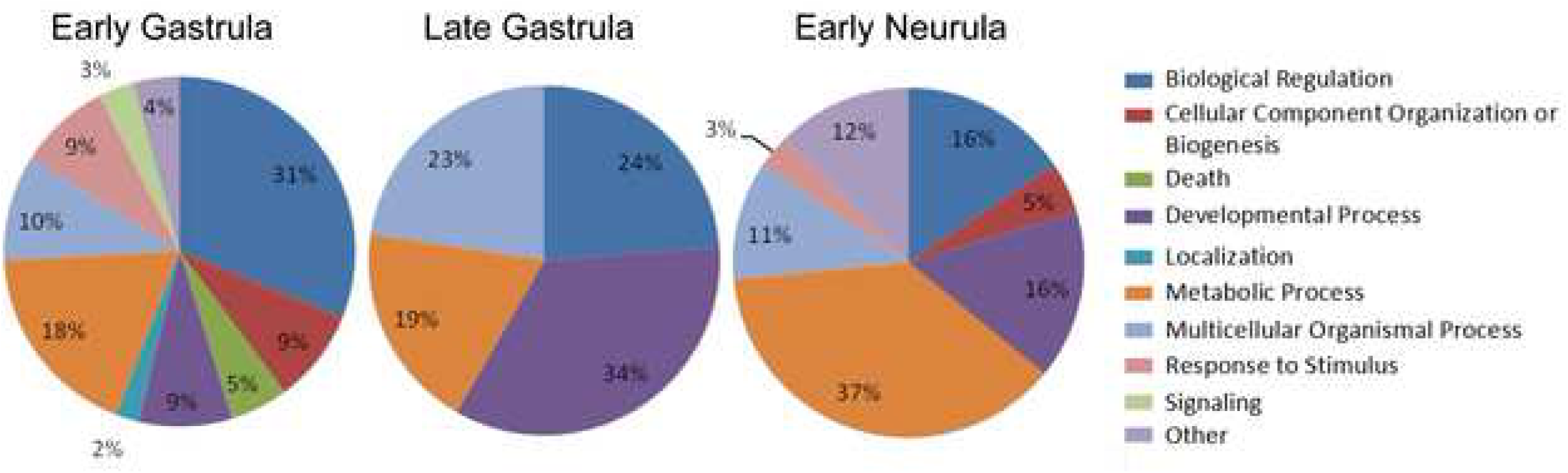
Biological processes most affected by loss of SUMOylation activity. Functional enrichment analysis of differentially expressed genes at each time point based on GeneGo categorization of annotated protein function. Only selected groups containing the largest number of genes are presented.

The expression of 9 out of 94 differentially expressed genes are assigned to Developmental Processes in the early gastrula embryos, of which the most notable are *foxc1, foxd1* and *twist*. All three of these genes, which are highly expressed in mesoderm, can be correlated with morphological phenotypes that later appear in the Gam1 embryos (*e.g.,* deficits in heart and neural tube development). Although GeneGo did not place *sox2*, a key regulator of neural induction, under Developmental Processes, it is nonetheless notable that its expression is reduced 1.7-fold at early gastrula and remains modestly lower (1.4-fold) at late gastrula.

The most striking change between early and late gastrula embryos is the greatly increased percentage of genes that fall under Developmental Processes (increasing from 9% to 34%) and Multicellular Organism Processes (increasing from 10% to 23%). Indeed, only four high level biological processes predominate at late gastrula. At this time point, several additional members of the Fox family of transcription factors appear in the list of differentially expressed genes ascribed to Developmental Processes as well as genes encoding proteins involved in Wnt, ephrin, and TGF-β signaling, which are all known to play critical roles during gastrulation.

Although the percentage of genes ascribed to Development Processes decreases in early neurula embryos to 16% of the total of differentially expressed genes, the actual number increases, showing that SUMOylation continues to play an important regulatory role in development. While the genes for many of the aforementioned transcription factors and signaling molecules remain mis-regulated, now multiple Hox genes are repressed in the Gam1 embryos at this time point, which is consistent with their normally strong induction during late gastrula stage. Whereas, SUMOylation is often associated with transcriptional repression, several of the Hox genes seemingly require this post-translational modification for activation.

Of special note, expression of the homeobox protein, Pitx2, is down 4-fold at early neurula in Gam1 embryos. This transcription factor is critical for eye development, as well as the formation of asymmetric organs such as the heart and gut, whose perturbation is a predominant phenotype of the SUMOylation deficient embryos. *Pitx2* along with *foxc1* (down 2.7-fold at early gastrula) are both implicated in Axenfeld-Rieger syndrome. The lists of genes that are categorized into the various biological processes can be found in Additional file 4: Table S1.

### Transcription factor network building

The regulation of transcription by SUMOylation can occur on multiple levels that include, for example, changes in chromatin structure (*e.g.,* histone modification and nucleosome positioning) and subnuclear localization (*e.g.*, PML bodies). However, the most common mechanism is likely through the SUMOylation of transcription factors. We used the network building algorithm ’Transcription Factor Regulation’ to identify common transcription factors that control the maximum number of differentially expressed genes in Gam1 embryos. This algorithm adds a transcription factor to a gene list as a seed node, and builds a network of interactions around it. Gene lists from each time point were analyzed separately and the top 30 transcription factors at each time point were returned (Table 1). The connections made within each network are based on direct control of the target gene by the seed transcription factor and, inversely, target gene control of the seed transcription factor. The lists of transcription factors at each time point are quite similar and include those such as Sp1, HNF4-α, c-myc, and p53, that regulate an especially large number of different genes.

**Table 1.** Transcription Factor Network Building^*^

| <u>Differential Gene Expression</u> |  |  |  |  |  | <u>Constant Gene Expression</u> |  |  |  |  |  |
| --- | --- | --- | --- | --- | --- | --- | --- | --- | --- | --- | --- |
| Early Gastrula<br>(94) |  | Late Gastrula<br>(447) |  | Early Neurula<br>(742) |  | Early Gastrula<br>(11,702) |  | Late Gastrula<br>(11,322) |  | Early Neurula<br>(11,019) |  |
| SP1 | 15 | HNF4α | 104 | SP1 | 119 | HNF4α | 1372 | HNF4α | 1335 | HNF4α | 1313 |
| HNF4α | 14 | SP1 | 101 | HNF4α | 114 | SP1 | 1082 | SP1 | 1053 | SP1 | 1028 |
| c-Myc | 13 | c-Myc | 92 | c-Myc | 106 | c-Myc | 1033 | c-Myc | 1002 | c-Myc | 987 |
| p53 | 9 | ESR1 | 75 | p53 | 77 | p53 | 486 | p53 | 461 | p53 | 460 |
| Oct-3/4 | 7 | p53 | 67 | ESR1 | 69 | ESR1 | 479 | ESR1 | 463 | ESR1 | 455 |
| STAT3 | 6 | E2F1 | 60 | C/EBPβ | 50 | CREB1 | 467 | CREB1 | 452 | CREB1 | 451 |
| CREB1 | 6 | NF-Y | 59 | NF-kB | 45 | AP-1 | 378 | AP-1 | 370 | AP-1 | 365 |
| YY1 | 5 | AR | 58 | STAT1 | 38 | NF-Y | 310 | E2F1 | 335 | E2F1 | 332 |
| HNF1α | 4 | EGR1 | 54 | AR | 37 | E2F1 | 342 | NF-Y | 301 | NF-Y | 303 |
| Sry | 4 | c-Jun | 53 | CREB1 | 37 | NF-kB | 314 | NF-kB | 302 | NF-kB | 298 |
| FKHR | 4 | ETS1 | 53 | HSF1 | 36 | AP-2 | 306 | AP-2 | 300 | YY1 | 294 |
| SRF | 4 | GCRα | 53 | MYOD | 36 | YY1 | 303 | EGR1 | 293 | AP-2 | 293 |
| NANOG | 4 | YY1 | 52 | SRF | 36 | EGR1 | 300 | YY1 | 290 | EGR1 | 289 |
| E2F1 | 4 | Oct-3/4 | 51 | EGR1 | 36 | GATA-1 | 283 | GATA-1 | 276 | GATA-1 | 266 |
| ETS1 | 4 | RelA | 51 | E2F1 | 36 | Elk-1 | 214 | Elk-1 | 211 | Elk-1 | 206 |
| EGR1 | 4 | Oct-1 | 50 | AP-1 | 35 | ETS1 | 250 | ETS1 | 248 | ETS1 | 242 |
| HIF1A | 4 | SRF | 50 | RelA | 34 | AR | 258 | AR | 255 | AR | 249 |
| AP-2 | 3 | HIF1A | 50 | C/EBPα | 32 | HIF1A | 236 | HIF1A | 233 | Oct-3/4 | 232 |
| ATF-6 $\alpha$ | 3 | MYOD | 49 | p21 | 32 | C/EBPβ | 235 | C/EBPβ | 230 | c-Jun | 224 |
| RelA | 3 | C/EBPβ | 49 | ATF-2 | 31 | Oct-3/4 | 236 | E2F4 | 228 | E2F4 | 223 |
| C/EBPα | 3 | Bcl-6 | 48 | YY1 | 29 | c-Jun | 233 | Oct-3/4 | 227 | HIF1A | 222 |
| ER81 | 3 | TCF7L2 | 48 | HIF1A | 29 | E2F4 | 229 | c-Jun | 222 | C/EBPβ | 217 |
| HSF1 | 3 | HSF1 | 48 | p63 | 29 | RelA | 224 | RelA | 216 | RelA | 214 |
| ESR1 | 3 | HNF6 | 48 | SP3 | 28 | SP3 | 206 | SP3 | 203 | SP3 | 199 |
| USF2 | 3 | E2F4 | 47 | NF-Y | 28 | GCRα | 205 | SRF | 188 | GCRα | 198 |
| AP-1 | 3 | SMAD3 | 47 | AP-4 | 28 | HNF6 | 205 | AP-2A | 197 | HNF6 | 198 |
| AR | 3 | SOX4 | 47 | c-Jun | 27 | Oct-1 | 202 | HNF6 | 197 | AP-2A | 194 |
| MYOD | 3 | STAT1 | 46 | GATA-1 | 26 | AP-2A | 200 | Oct-1 | 195 | Oct-1 | 192 |
| FOXO3A | 3 | c-Myb | 46 | STAT3 | 26 | AHR | 192 | GCRα | 195 | HNF1α | 187 |
| AP-2A | 3 | SREBP1 | 46 | Oct-3/4 | 24 | HNF1α | 192 | AHR | 189 | SRF | 185 |

\*Top 30 transcription factors associated the largest number of genes that are either differentially expressed or constant across developmental stages of Gam1 embryos built using the Transcription Factor Regulation algorithm in MetaCore. Numbers indicate the number of genes used for network building in each case.

In order to determine if these transcription factors are highly associated with genes controlled by SUMOylation, the same analysis was carried out using the lists of genes not affected by Gam1, which number 11,702 genes from early gastrula, 11,322 genes from late gastrula, and 11,019 genes from early neurula (Table 1). The list of transcription factors connected to changed *versus* unchanged genes is strikingly similar. At early gastrula, 18 out of 30 transcription factors were common to both sets of genes with the top 5 factors being identical. For the late gastrula, 21 out of 30 factors are common and again the top 5 are identical. At the early neurula, 20 out of 30 factors are common with the top 5 factors remaining identical between the lists. Of the top 5 transcription factors, four are common across all three time points (SP1, HNF4-α, c-myc, p53). The simplest interpretation of this analysis is that these transcription factors emerge solely because they bind to an unusually large number of regulatory sites throughout the genome, a subset of which are regulated by SUMOylation.

Interestingly, the GO biological processes associated with any specific transcription factor are different depending on whether the gene list is built from those changed by Gam1 or those unaffected by Gam1 (Additional file 5: Table S2). For example, the two top transcription factors at early gastrula are Sp1 and HNF4-α. The GO processes most highly associated with Sp1 in Gam1 embryos are eye, sensory organ, and embryo development; whereas, in control embryos they are metabolic processes. It is especially noteworthy that SUMOylation of Sp1 has been directly implicated in *Xenopus* eye development [37]. For HNF4-α, the enriched GO processes in Gam1 embryos are dendrite morphogenesis, signaling pathways, microtubule polymerization, and cell communication; the GO processes in control embryos are almost entirely metabolic. This analysis indicates that constitutive genes necessary for organism viability, *e.g.,* metabolism and biosynthesis, are mostly controlled in a SUMO-independent manner. Conversely, genes with complex expression patterns are more likely controlled by transcription factors whose activity is modulated by SUMOylation. This GO analysis also suggests that SUMO modification generates distinct pools of a particular transcription factor in order to regulate specific subsets of genes, with SUMOylated forms of these transcription factors biased towards more highly regulated (*e.g.,* cell type, temporal) processes. This view is supported by studies in *Drosophila* where mutations in the SUMO E1 activating enzyme had no effect on larval growth, but did cause defects in imaginal disk formation [38].

A total of 50 different transcription factors were identified through the Transcription Factor Regulation algorithm as being associated with genes differentially expressed at one or more time points. These factors were analyzed for potential SUMOylation sites and SIMs using the SUMO prediction site algorithm GPS-SUMO [39]. Of the 50 transcription factors, 46 (92%) contain consensus SUMOylation sites, only 1 has no identifiable site, and 3 lack sufficient sequence information to make a prediction (Additional file 6: Table S3). To date, 29 of the 46 have been experimentally verified to be SUMOylated. SIMs appear in 42 of the listed transcription factors. The fact that nearly all of factors identified by network building are targets of SUMOylation and/or contain SIMs indicates that the majority of changes in gene expression induced by Gam1 are due directly to changes in factor activity as a result of this post-translational modification.

The Transcription Factor Regulation algorithm returns the most highly connected factors for a given gene list. A shortcoming of this analysis is that critical transcription factors may be overlooked, if they regulate a small fraction of genes in the entire list. As an example, the gene list at early gastrula was examined manually using a variety of sources to identify candidate transcription factors for each gene. As expected, we found that the transcription factors listed in Table 1 are associated with many of the differentially expressed genes in the list. However, we also found that the transcription factor Nkx2.5, whose role in cardiac differentiation is regulated by SUMOylation [28], has predicted binding sites at the promoters of a subset of these differentially expressed genes (*alkbh6, evx1, fam184a, foxc1, foxd1, fzd10a, gapvd1, twist1/2*). Of these putative targets of Nkx2.5, *foxc1* is particularly significant, since *foxc1*-depleted *Xenopus* embryos exhibit similar phenotypes to the Gam1-injected embryos: shortened A-P axis and abnormal heart and gut development [40]. Moreover, a morpholino knockdown of foxc1 protein caused changes in the expression levels of *casein kinase 1 epsilon*, *insulin-like growth factor 2*, and *keratin 8* mRNAs [40] that we also detect in the early neurula embryos.

The heart defects of Gam1-injected embryos are certainly not due to *foxc1* alone. We also measured reductions of other key cardiac regulatory factors such as *mespa* (1.6-fold), *tbx2* (2.1-fold), and *tbx3* (2.3-fold) at late gastrula and *pitx2* (4.0-fold) at early neurula. Thus, the potential effects of depleted SUMOylation activity on heart development are seemingly propagated through the three stages sampled in this study. *Twist,* which is down regulated at early gastrula and considerably more at late gastrula, has also been implicated in cardiogenesis (reviewed in [41]). Thus, the predominant phenotypes in heart development can be explained by multiple effects on the cardiac transcription program that results from depletion of SUMOylation activity.

### Co-expression analysis

The expression of the differentially expressed genes were compiled across the three time points and compared using the module/cluster detection algorithm implemented in R package WGCNA [42]. The differentially regulated genes were compiled into eight syn-expression clusters based on the similarity of expression changes across all three time points. The gene lists from individual clusters were analyzed using the MataCore transcriptional regulation algorithm. Each cluster returned a similar set of top transcription factors that are similar to the individual lists returned for both the differentially expressed and unchanged genes (Table 1). This observation further supports the idea that the most highly connected transcription factors are those that are widely used to control the greatest number of genes in any list. This strongly argues that co-expression in this case cannot be traced to a single transcription factor or set of transcription factors acting as master regulators.

### Signaling pathways sensitive to diminished SUMOylation activity

We chose a non-lethal level of Gam1 expression in order to identify developmental processes that are potentially regulated by SUMOylation. This strategy will only identify those that are most sensitive to loss of this activity. Because the effects of this post-translational modification are widespread and not limited to gene regulatory proteins, we have used the predominant phenotypes of Gam1-injected embryos as a guide to examine some of the signaling pathways that appear to be the most vulnerable to this perturbation. The list of differentially expressed genes was uploaded into MetaCore and analyzed for their presence in known biochemical pathways. Analysis focused on those pathways with outcomes related to the observed phenotypes of failed blastopore or neural tube closure, shortened A-P axis, and cardiovascular deformations. Three pathways, (i) non-canonical Wnt signaling, (ii) epithelial to mesenchymal transition (EMT), and (iii) the Ets-1 pathway, contain multiple differentially expressed genes and were chosen as examples of cases where disruption of SUMOylation can be related to the observed phenotypes.

### Non-canonical Wnt/PCP signaling

Wnt signaling can be divided into two broad groups, canonical and non-canonical, based on outcome. Canonical signaling is dependent on β-catenin and regulates transcription of target genes, while non-canonical is independent of β-catenin and regulates cell polarity and calcium levels [43]. The non-canonical signaling pathway leads to activation of Rho and reorganization of the cytoskeleton and changes in cell polarity. Specific Wnt proteins are classified as canonical or non-canonical based on their ability to induce secondary axes in *Xenopus* embryos. The latter, which includes Wnt4, Wnt5a, and Wnt11 [44–46] have little or no dorsalizing effect and do not activate known Wnt target genes such as *nodal3.1* (*Xnr3*) and *sia1* (*Siamois*), whose expression is not changed in Gam1 embryos.

Several Wnt and Fzd genes are misregulated in SUMOylation deficient embryos (Fig. 5). Three Fzd genes of the non-canonical pathway (*fzd2, fzd7* and *fzd10*) are down regulated at one or more time points. Two Wnt genes (*wnt8b* and *wnt11*) are also down regulated. Wnt8b has not been specifically implicated in non-canonical signaling; however, it is often placed in the same group as Wnt11, since both have similar temporal roles, activating canonical pathways in oocytes and pre-MBT embryos, but non-canonical pathways during gastrulation [47, 48]. Both Wnt8 and Wnt11 can bind to Fzd7 [49].

**Figure 5.**
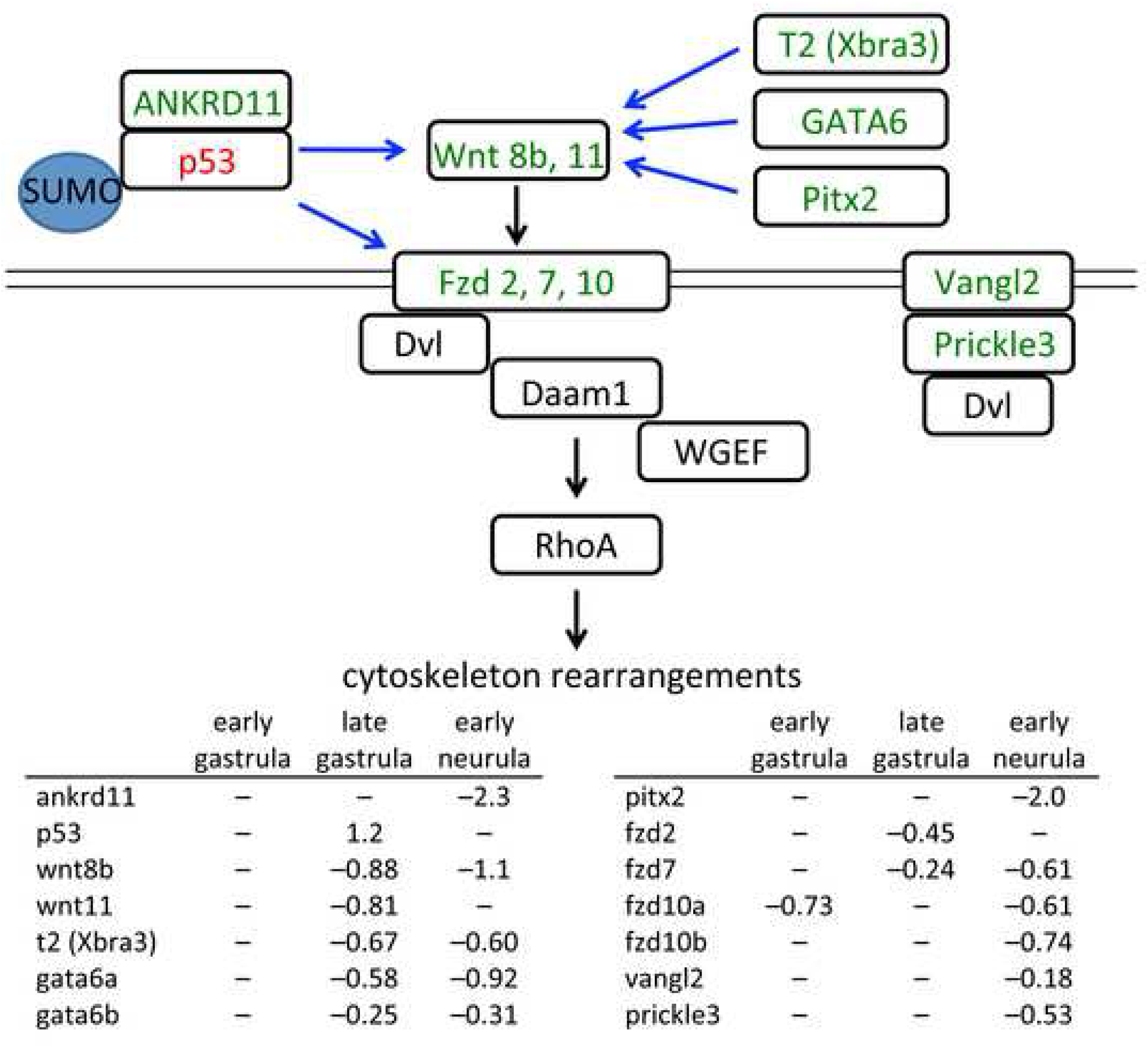
Changes in gene expression in the non-canonical Wnt/PCP signaling pathway. Decreased mRNA levels in Gam1 embryos relative to control embryos is represented by *green* and increased levels by *red*. Documented targets of SUMOylation are indicated. Transcriptional regulation is denoted by *blue* arrows and signaling through protein-protein interactions by *black* arrows. Log(2) changes in gene expression are listed for the three experimental time points.

While Fzd2 and Fzd7 have been implicated in planar cell polarity (PCP), the interaction of the latter receptor with Wnt11 ligand appears to be the major determinant of this process [50–53]. Therefore, the observed decrease in both these receptors and ligands can account for the effect of Gam1 on convergent extension. An early manifestation of convergent extension/PCP is closure of the blastopore, which is generally delayed and often incomplete in Gam1 embryos (Fig. 2A) and is specifically attributed to forces generated by convergent thickening (CT) [54]. Convergent extension underlies the intercalation of mesodermal cells that drives the elongation of the A-P axis, which is also compromised in the SUMOylation deficient embryos. Both Wnt 11 and Fzd7 have been implicated in A-P axis elongation. A constitutively active form of the Rho GTPase, Cdc42, rescues axis elongation effects caused by expression of a truncated form of Fzd7, while a dominant negative form of Cdc42 rescues the inhibition of axis elongation that occurs with Wnt11 and Fzd7 overexpression [50]. In some cases, Fzd7 and Fzd2 act redundantly in convergent extension [55], which likely accounts for the high penetrance of convergent extension phenotypes in Gam1 embryos, since we measure a decrease in expression of both receptors during late gastrula.

Disruption of the non-canonical Wnt pathway can also account for the cardiac and neural tube defects that occur later in embryo development. Wnt 11 and Fzd7 are required at multiple points during *Xenopus* heart development, not only early in the specification of cardiac progenitor cells, but also in cardiac morphogenesis (septation and outflow tract development) [56–61]. The ensuing decreased activity of Dishevelled (Dvl) in Gam1 embryos is fully consistent with the cardiac and neural tube phenotypes of Dvl-deficient mice and *Xenopus* [62–64].

Two other PCP core components, Vangl2 and Prickle3, are modestly reduced at early neurula stage, a period when a complex of these two proteins is required for the radial cell intercalations that sustain neural tube closure [65, 66]. Notably, the polarization of the Prickle3/Vangl2 complex to anterior cell edges has been linked to Wnt11 signaling [67]. Vangl2 also plays a critical role in heart formation that is independent of neural tube closure. In mice, knock down of Vangl2 specifically in undifferentiated cells of the second heart field prevents lengthening of the outflow tract due to a failure of these cells to polarize and differentiate [68]. These early abnormalities lead to later defects in cardiac morphology. Decreased expression of several members of the non-canonical Wnt signaling pathway combined with those of the PCP pathway can account for the most prominent phenotypes of the SUMOylation-deficient *Xenopus* embryos and strongly suggest that disruptions in this post-translational modification potentially underlie congenital birth defects of the neural tube and heart [30, 69].

Because several Wnt and Fzd genes are down regulated in the Gam1 embryos, it is reasonable to speculate that they share one or more common transcriptional regulators that can coordinate their expression; however, it has not been possible to discern a single master regulator. Transcription factor network building indicated p53 activity as the most likely source for dysregulation of non-canonical Wnt signaling. At all three time points, p53 is in the top five transcription factors most highly connected to differentially expressed genes. Surprisingly, p53 shows a two-fold increase in expression at late gastrula in Gam1 embryos. However, regulation of p53 activity by SUMOylation is complex with the modification having a positive, negative, or no effect depending on the target gene in question [70]. Moreover, ANKRD11, a coactivator of p53 [71], shows a five-fold decrease in expression by early neurula stage. Our results indicate that SUMOylation of p53 and/or one of its partners plays a critical role in the regulation of non-canonical Wnt pathway genes during *Xenopus* gastrulation and neurulation (Fig. 5). In addition, decreased expression of transcription factors GATA6 [72], Pitx2 [73], and T2 (Xbra3) [74] can also contribute specifically to the decrease in *wnt11* mRNA expression (Fig. 5).

### Epithelial to mesenchymal transition

The epithelial to mesenchymal transition (EMT) is a hallmark of early vertebrate development and patterning of the embryo [75, 76]. In *Xenopus*, EMT changes allow for the presumptive mesoderm to migrate inward during gastrulation, and disruptions in EMT signaling lead to developmental defects in gastrulation and later patterning of the embryo [77]. One important pathway involves control of EMT genes by the transcription factors snail 1 (*snai1*) and twist (*twist1*) [75, 76, 78]. Although the expression of *snai1* and *twist1* genes requires high mobility group A-T hook 2 (HMGA2), the two genes appear to be under distinct regulatory control [79]. HMGA2 binding to the *snai1* promoter involves formation of a complex with phosphorylated SMADs activated by TGFβ signaling [80]. Conversely, HMGA2 is able to bind directly to the *twist1* promoter in the absence of TGFβ signaling [79, 80]. HMGA2 is a known target of SUMOylation [81, 82] that has been directly implicated in *Xenopus* EMT and migration of neural crest cells [83] as well as *Xenopus* cardiogenesis [84]. The different behavior of HMGA2 at the *snai1* and *twist1* promoters is reflected in the different effect of Gam1 on the expression of these two genes. *Twist* expression was reduced at all three time points. In contrast, no misregulation of *snai1* was observed.

The microarray analysis detected disruptions in the twist/snail pathway that potentially increase epithelial and decrease mesenchymal characteristics that account for the observed gastrulation defects and failure of the blastopore to close (Fig. 6). Twist and Snail1 form a complex to control expression of the EMT-specific transcription factor, Zeb-2, which is down regulated 1.3-fold at early neurula. Zeb-2 is responsible both for activation of genes that promote a mesenchymal phenotype and repression of genes that promote an epithelial phenotype [85]. Moreover, Zeb-2 is a target of SUMOylation, which controls its activity in a promoter-specific fashion [86]. ZO-2 (*tjp2*), an integral component of tight junctions normally down-regulated by Zeb-2 in mesenchymal cells, allowing for increased cell movement, is up-regulated at late gastrula and early neurula in Gam1 embryos. Vimentin (*vim*), claudin 4 (*cldn4*), and connexin26 (*gjb2*) normally up regulated in mesenchymal cells, are down regulated in Gam1 embryos [87, 88].

**Figure 6.**
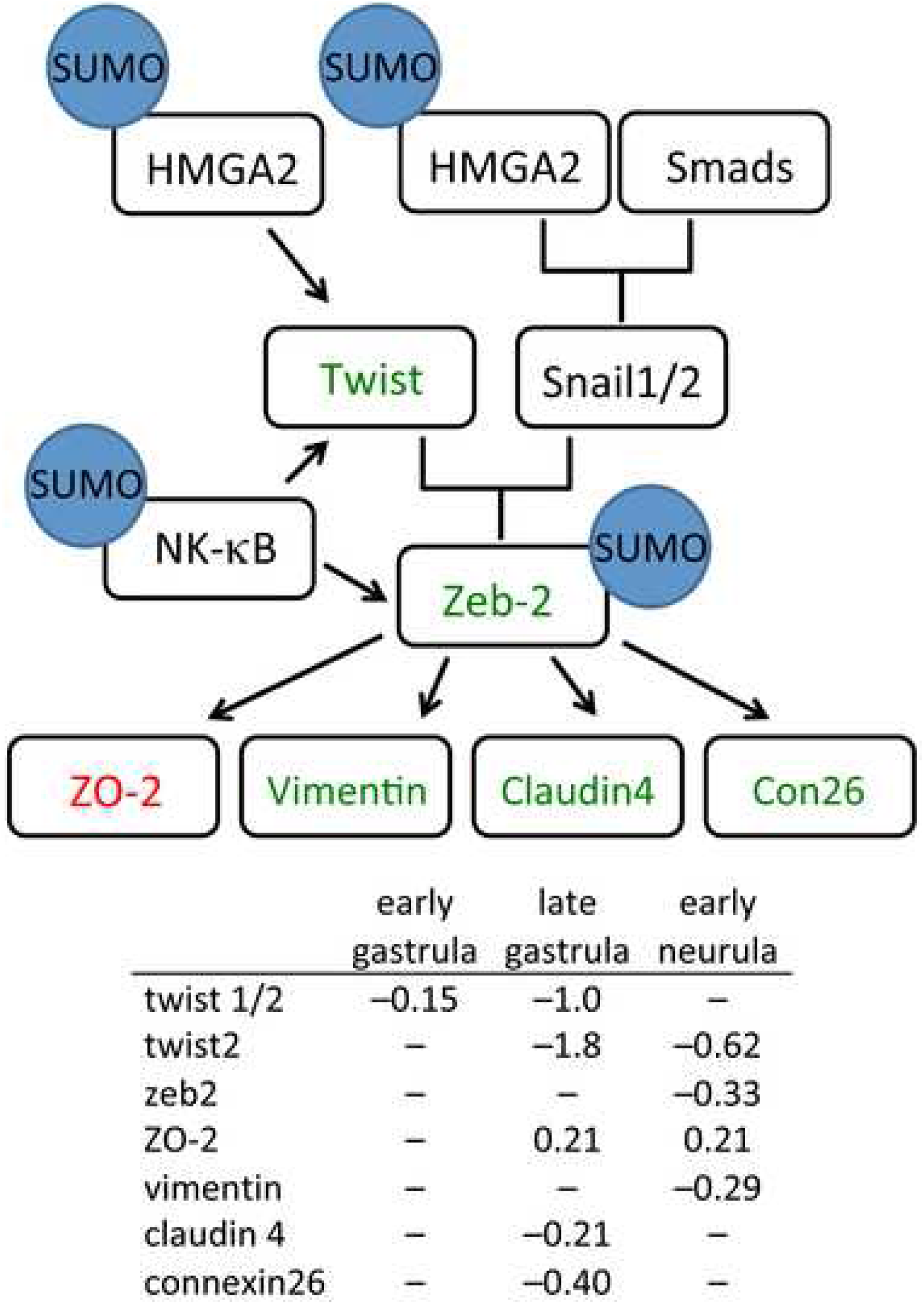
Changes in gene expression in the Twist/Snail pathway. Decreased mRNA levels in Gam1 embryos relative to control embryos is represented by *green* and increased levels by *red*. Documented targets of SUMOylation are indicated. Transcriptional regulation is denoted by *black* arrows. Log(2) changes in gene expression are listed for the three experimental time points.

SUMO possibly impacts this pathway at a second point. NF-κB/p65 an activator of Twist and Zeb2 expression [89], is regulated by a negative feedback loop involving the SUMOylation of the p65 (RELA) subunit [90] that can account for the decreased expression these two transcription factors. In combination, the altered expression of the proteins in this pathway is expected to contribute to an epithelial-like character that impedes cell migration during gastrulation and neurulation, resulting in phenotypes such as the observed incomplete closure of the neural tube and shortened A-P axis. On the other hand, the constant level of *snail1* expression can account for the unperturbed levels that we measure of other regulators of the EMT (*e.g.,* E-cadherin, N-cadherin, fibronectin, and occludin).

### Ets-1 pathway

Pathway analysis of the microarray data revealed that perturbation of Ets-1 activity is another likely contributor to the phenotypes exhibited by Gam1 embryos [91–94]. The regulation of Ets-1 during development is complex and occurs through the integration of multiple types of post-translational modification: phosphorylation, acetylation, ubiquitination, and SUMOylation [92, 94, 95]. Furthermore, Ets factors can either enhance or repress transcription [96]. Ets-1 can be SUMOylated at two lysine residues (K15 and K227), resulting in the loss of transcriptional activation activity [92]. Conversely, Ets-1 phosphorylated at threonine 38 by MAPK, as a result of Ras activation, increases its transcriptional activation activity (Fig. 7). The SUMOylation of two targets in the Ras/Raf-MEK-MAPK phosphorylation pathway indirectly influence Ets-1. Ras-1 is a target for SUMO modification, which is required for Ras-dependent activation of MAPK [97]. In contrast, SUMOylation of the downstream target MEK blocks its interaction and activation of MAPK [98]. Down regulation of several genes controlled by Ets-1 in Gam1 embryos indicates that inactivation of the Ras pathway and the resulting dephosphorylation of Ets-1 has the greatest impact on the activity of this transcription factor.

**Figure 7.**
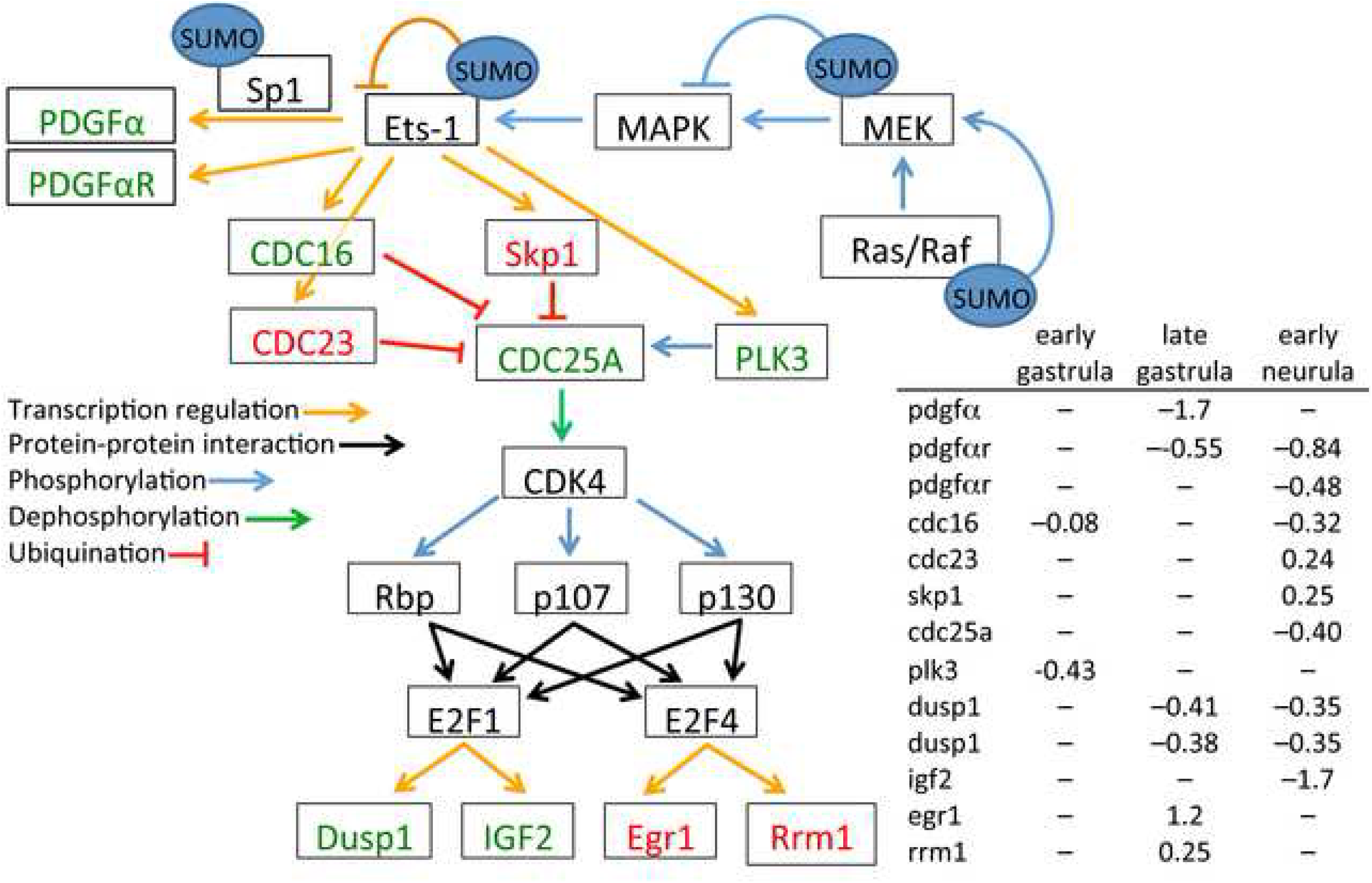
Changes in gene expression in the Ets-1 pathway. Decreased mRNA levels in Gam1 embryos relative to control embryos is represented by *green* and increased levels by *red*. Documented targets of SUMOylation are indicated. Connections by arrows represent activation and T-bars represent inhibition. Log(2) changes in gene expression are listed for the three experimental time points.

Knockdown of Ets-1 in *Xenopus* neural crest cells results in defects in delamination of the neural tube and disruption of cell migration that lead to malformed outflow tracts of the heart; whereas, depletion of Ets-1 activity in heart mesoderm results in distinct phenotypes that include loss of the entire endocardium and failure to form discrete chambers [93]. Two Ets-1 target genes, platelet derived growth factor α (*pdgfa*) and PDGFα receptor (*pdgfra*), are down regulated. There is abundant evidence that PDGFα is necessary very early in heart development for the proper migration of cardiomyocytes to the midline and formation of the heart tube [99–101]. Ets-1 works in concert with Sp1 to activate transcription of the PDGFα gene [102]. Like Ets-1, Sp1 is repressed by SUMOylation [103] and activated by MAPK [104], potentially amplifying the effect of Gam1 on the expression of *pdgfa* and *pdgfra*.

Gastrulation defects in developing *Xenopus* embryos are also observed when PDGFa signaling is disrupted [105–107]. PDGFa is expressed in the blastocoel roof of the developing embryo and its receptor is expressed in the migrating mesoderm. This complementary expression of ligand and receptor orients and directs the movement of mesodermal cells inward during gastrulation. Interfering with PDGFa signaling randomizes internal movement of these cells and prevents proper convergent extension [106]. This disruption leads to improper patterning and phenotypes that include incomplete blastopore closure and shortened axis elongation. Mice with mutations in the PDGFaR display severe skeletal abnormalities, cleft face, and *spina bifida* [108, 109], with the latter being a frequent phenotype in Gam1 embryos.

While the disruption of Ets-1 (and Sp1) activity can account for the immediate effects on PDGFα and its receptor, we note that there are also several downstream changes in gene expression that can contribute to the phenotypes of Gam1 embryos, including DUSP1 (cell migration) [110] and IGF2 (neural and eye development) [111, 112]. Of particular note, is the increased expression of the transcription factor Egr1 (late gastrula) which is a positive regulator of MyoD and a negative regulator of Xbra [113]. Consistent with the change in Egr1, we measure increased expression of MyoD (late neurula) and decreased Xbra expression (late gastrula/early neurula). Both MyoD and Xbra are critical factors in several developmental pathways.

## Conclusions

It is now appreciated that SUMO impinges on an substantial number of biological processes, explaining why deletion of E2 conjugating enzyme is embryonic lethal in mice and zebrafish [2, 8, 11–13]. For this reason, it has been difficult to assess the role of this post-translational modification in live animals. While the magnitude of many of the changes in gene expression in Gam1 embryos is modest, the cumulative effect of disrupting multiple pathways that impinge upon a particular developmental event accounts for the frequency of the commonly observed phenotypes. The experiments here make evident that regulation by SUMO during the earliest period of embryogenesis is critical to proper development at later stages. Injection of Gam1 mRNA suppressed SUMOylation activity up to late neurula (26 hpf), yet generated phenotypes that appear later in development (*e.g.,* heart looping, 48 hpf), demonstrating the early embryonic origins that give rise to the two most common human birth defects: heart and neural tube. Transcriptome analysis has revealed those pathways that are most susceptible to diminished SUMOylation activity. The frequent occurrence of heart defects in Gam1 embryos is in accord with the prominent role SUMO plays in the regulation of cardiac gene expression [114, 115]. Indeed, in a cohort of 87 babies exhibiting atrial septal defects, five carried mutations that reduced the promoter activity of the SUMO1 gene [30], demonstrating a direct connection between deficits in this posttranslational modification and congenital heart defects. It is also clear from the experiments described here that the PCP pathway is exceptionally sensitive to depressed SUMOylation activity and this most certainly accounts for the frequent neural tube defects of the Gam1 embryos [116, 117] and supports a proposed relationship between the PCP pathway and risk for spina bifida [69]. Our transcriptomics analysis has identified disruptions in pathways that have especially great potential to contribute to these pathologies and that warrant closer clinical examination. Targeted inhibition of SUMOylation activity by Gam1 will be useful for further investigation of this post-translational modification and its causative role in birth defects.

## Methods

### Embryo preparation

Female *X. laevis* were injected with 500 units of human chorionic gonadotropin (HCG) at least 12 hours prior to spawning. Testes were isolated and stored at 4°C in high salt MBS (0.7 mM CaCl_2_, 108 mM NaCl, 1 mM KCl, 1 mM MgSO_4_, 5 mM HEPES, pH 7.8, 2.5 mM NaHCO_3_) for up to a week. Minced testis and eggs were mixed together with 3 mL 1/3 MMR (0.1 M NaCl, 2.0 mM KCl, 1 mM MgSO_4_, 2 mM CaCl_2_, 5 mM HEPES, pH7.8) and left to fertilize at room temperature. After 10 minutes, eggs were flooded with additional 1/3 MMR and after an additional 30 minutes eggs were washed with 2% cysteine (< 5 minutes) in order to remove the jelly coating, upon which eggs were washed 8 times with 1/3 MMR. Embryos were injected in the animal hemisphere (volumes ranged between 5 to 30 nl) and allowed to develop at room temperature in 1/3 MMR. At least 30 minutes prior to harvest of the testes, the male frog is submerged in an ice water bath using a mesh basket to keep the ice away from the skin surface. The frog is decapitated, and pithed prior to removal of the testes. After removal of the testes tissue, the heart is cut and the frog pithed to ensure death prior to disposal. Female frogs are rested at least 3 months between spawnings.

### RNA synthesis

Linearized plasmid (2.5 ug) was added to a mixture containing transcription buffer (Promega), 10 mM DTT, 2.5 mM cap analog, 50 units RNasin, 200 units RNA polymerase, and 0.5 mM each UTP, ATP, CTP, GTP in a total volume of 50 uL. The reaction was incubated at 37 °C for 90 minute, followed by the addition of another 200 units of polymerase and incubation for 90 additional minutes. The reaction mixture was extracted twice with phenol pH 4.5 and twice with 24:1 chloroform: isoamylalcohol. The mRNA was precipitated with ethanol, suspended in 50 ul of H_2_O and run though a NucAway spin column (Ambion) to remove unincorporated nucleotides. mRNA concentrations were determined using a NanoDrop spectrometer (Thermo Scientific), and samples were stored at -80 °C until use.

### SUMOylation Assays

Gam1 mRNA was injected into one-cell embryos, which were allowed to develop to the indicated stage (midblastula, late blastula, early gastrula, and midneurula). Twenty embryos per time point were homogenized in 22 uL of SUMO reaction buffer (Boston Biochem) and spun at 16,000 rcf for 5 minutes. Supernatant was removed and the protein concentration was determined by Bradford assay. Each assay conatined 25 ug of whole cell extract, 60 uM SUMO1, 5 uM Ubc9, 5 uM E2-25K (substrate peptide), 25 mM Mg-ATP in a total reaction volume of 25 uL. Control reactions contained 500 nM SAE1/SAE2 (E1 enzyme) in place of cell extract. Reactions were incubated at 37 °C for four hours. SDS loading dye was mixed with the reactions and vortexed briefly in order to stop the reaction. Assays were analyzed by SDS polyacrylamide gel electrophoresis followed by western blot.

### Western blots

Gam1 carries a single (10 amino acid) N-terminal myc tag that was detected using c-myc antibody (sc-40, Santa Cruz Biotechnology) at 1:500 dilution. For SUMOylation assays an antibody (A-603, Boston Biochem) directed against the SUMO1 substrate peptide (E-25K) was used at 1:500 dilution. All blots were visualized using a goat anti-rabbit IgG:alkaline phosphatase fusion protein at 1:3,000 dilution in conjunction with BCIP/NBT color development (Bio-Rad).

### Keller sandwich explant assays

One-cell embryos injected with either Gam1 mRNA (2.5 ng or 5.0 ng) or an equivalent volume of H_2_O and allowed to develop in 1/3 MMR until early gastrula stage (~9 hpf) when the blastopore lip was initially beginning to form. Embryos were maintained in 1× MBS buffer (88 mM NaCl, 1 mM KCl, 0.7 mM CaCl_2_, 1 mM MgSO_4_, 5 mM HEPES pH 7.8, 2.5 mM NaHCO_3_) during the dissection, which followed the procedure described, by Keller and Danilchik (1998).

Once the sandwich was fabricated, it was cultured in the dissection dish for one hour allowing the explants to adhere. Explant sandwiches were then moved to an agarose-coated Petri dish containing 1X MBS supplemented with 50 ug/mL gentamicin and left to culture for an additional 14 hours. Samples were photographed immediately after moving them to the Petri dish (10 hpf) and again at 24 hpf using an Olympus SZX16 microscope. Scale bars on the CellSens software were used to measure the length and width of the sandwich at the mentioned time points. Explants from water-injected or Gam1-injected embryos were compared to each other to determine the level of convergence and extension. Sample sets were analyzed for significant differences using a Student’s T-Test.

### Microarray analysis

One cell embryos were injected with either 0.5 ng Gam1 mRNA in a total volume of 9.2 nL or with H_2_O as a control. Embryos were allowed to develop in 1/3 MMR at room temperature until the desired time points. Samples were taken at early gastrula (9 hpf), late gastrula (13.5 hpf) and early neurula (16.5 hpf). Twenty embryos from each time point were homogenized in 500 uL proteinase K buffer (100 mM Tris pH 8, 150 mM NaCl, 12.5 mM EDTA pH 8.0, 1% SDS, 200 ug proteinase K) and incubated at 37 °C for one hour. Samples were then extracted twice with an equal volume of phenol, pH 4.5, and twice with 24:1 chloroform:isoamylalcohol. RNA was purified using an RNAEasy spin column (Qiagen, #74104) and precipitated with 2.5 volumes of ice-cold ethanol. The concentration and purity of the samples was determined using a NanoDrop spectrometer. Prior to microarray data collection, the RNA integrity number (RIN) for each sample was determined by the Notre Dame Genomics and Bioinformatics Core Facility (GBCF) using an Agilent 2100 BioAnalyzer.

Total RNA (50-500 ng) was used for cDNA synthesis according to the GeneChip 3’ IVT Express Kit for Affymetrix microarray chips. The cDNA was then converted to double-stranded DNA and used as a template for transcription of amplified RNA (aRNA) which was labeled with a biotin-conjugated nucleotide. aRNA samples (15 ug) were subsequently purified and fragmented to produce probes for hybridization onto *Xenopus laevis* 2.0 GeneChip 3’expression arrays. Three biological replicates were carried out for Gam1 and H_2_O injected samples at each time point utilizing a total of 18 microarray chips. Analysis of the microarray chips was carried out with the Affymetrix GeneChip System. Affymetrix .cel files were analyzed using the Bioconductor software package (http://www.bioconductor.org/). The mean fluorescence intensity was derived from a log_2_ transformation of the data and normalized using the quantile normalization method. A Student’s t test was used to determine if there was a significant difference between the Gam1 and H_2_O embryo gene expression values. A *p* value less than 0.05 was used as a cut off for differential expression. The fold change, between control and Gam1 samples, for each expression value was also calculated. Gene lists for those genes differentially expressed (p<0.05) were compiled for bioinformatics analysis. Since the *Xenopus laevis* genome is not fully annotated, some probes on the 2.0 chip, which do not have identified names or functions; only annotated genes were used for bioinformatics analysis.

### Microarray data analysis: MetaCore

Gene lists were uploaded to the MetaCore software suite of programs; only annotated genes were used for each analysis and genes were grouped differently depending on the analysis. Genes were grouped into lists based on differential expression at each time point (EG, LG, EN), not-differential expression at each time point (EG, LG, EN), or common expression patterns across all three time points (Cluster 1-8). MetaCore was used to analyze the gene lists for (i) transcription factor networks, (ii) functional enrichment, and (iii) pathway analysis each using an algorithm with specific well-defined instructions for creating the network based on established interactions.

In order to remove the redundancies created by the MetaCore enrichment algorithms, GeneGo IDs identified in the initial analysis were further categorized using GO Term classification software CateGOrizer (v. 3.218). Gene ontologies are hierarchically organized and the CateGOrizer software allows the specific sub-terms to be grouped into their broader categories providing a more accurate enrichment picture.

Pathway maps in MetaCore are created by compiling known biochemical processes or signaling cascades into graphical images. These maps can be used to determine the interconnectedness of uploaded genes in a biological context. Additionally, pathway maps can be used to determine downstream gene targets not present on the microarray. Gene lists were uploaded into MetaCore while pathways relevant to development (heart and nervous system), cell adhesion, cell cycle, cytoskeleton remodeling, and epithelial to mesenchymal transition were manually searched for any matching genes. Pathways containing differentially regulated genes were analyzed further for their connection to the observed developmental phenotypes.

### Identification of SUMO consensus sequences and SIMs

Potential sites of SUMOylation and SUMO interaction motifs (SIM) were identifed using GPS-SUMO (http://sumosp.biocuckoo.org) using the low threshold setting for the former. *X. laevis* reference sequences were accessed through XenBase (www.Xenbase.org)

### Real time polymerase chain reaction

RNA samples (4 ug) were mixed with random hexamers (Promega, #C118A) and heated at 70 °C for 5 minutes. Samples were then placed on ice for at least 5 minutes. Reverse transcription reactions containing 1 uL GoScript reverse transcriptase (Promega, #A501C) in a final volume of 20 uL were placed in a thermal cycler with program sequence: 25 °C for 10 min, 42 °C for 50 min, 70 °C for 15min. Dilution calculations of the reverse transcription (RT) reaction were based on a proportional amount of cDNA being created from the total RNA added to the reaction thereby producing a solution with a cDNA concentration of 200 ng/uL. RT samples were diluted with H_2_O to give a final cDNA concentration of 5 ng/uL. An aliquot of 50 uL of cDNA was mixed with an additional 200 uL of H_2_O to give a final working concentration of 1 ng/uL which was used in SYBR green PCR reactions. An additional 50 uL aliquot of each 5 ug/uL RT sample was taken in order to make standard curve dilutions for testing primer efficiency. A four point standard curve was created with 10 fold dilutions from the 5 ng/uL cDNA arbitrarily termed: 1000 (undiluted), 100, 10, and 1. A standard curve was generated for each primer pair in order to test the primer efficiency. A master mix containing 25 uM of both forward and reverse primers (Table 1) and 12.5 uL SYBR Green Master Mix (Applied Biosystems, #4309155) was prepared for each sample. The master mix (15 uL) and cDNA (10 ul) was added to each well of a 96 well plate. Samples were run with the following program parameters: 50 °C for 2 minutes, 95 °C for 15 minutes, 95 °C for 15 seconds, 60 °C for 30 seconds, 72 °C for 30 seconds (last three steps repeated for 45 cycles), 95 °C for 15 seconds, 60 °C for 1 minute, ramping temperature for 20 minutes, 95 °C for 15 seconds. The ramping temperature function was used to produce melt curves for calculation of primer efficiency.

## Abbreviations

Dvl: Dishevelled
GO: gene ontology
MBT: midblastula transition
PCP: planar cell polarity
SIM: SUMO interaction motif

## Declarations

### Ethics and Consent

Not applicable

### Research Involving Animals

All animal experiments were performed in accordance with protocols approved by the University of Notre Dame Institutional Animal Care and Use Committee (Protocol Number 18-02-4408).

### Consent for Publication

Not applicable

### Trial Registration

Not applicable

### Availability of Data and Materials

All data files have been uploaded to the Gene Expression Omnibus repository, including, experiment protocols, sample specifications, original CEL files, and processed Log2 GC-RMA signals under the following accession number GSE116164. Plasmids used in this study are available upon request from the corresponding author.

### Author’s contributions

MMB and PWH conceived and designed the project. MMB performed the experiments. KMD performed the Gam1 western blot. MMB, LC, EZ, and PWH carried out the bioinformatics analysis. MMB, EZ, and PWH wrote the manuscript with input from all authors.

## Acknowledgments

We thank Dr. Olivia Cox for assistance with the bioinformatics analysis and members of the Notre Dame Genomics & Bioinformatics Core Facility. We are grateful to Dr. Susanna Chiocca for providing a clone of Gam1. We acknowledge the expertise acquired from the organizers and instructors of the Cold Spring Harbor Laboratory *Xenopus* course. Constructive comments on the manuscript came from Olivia Cox, Joseph O’Tousa, Holly Goodson, and Kyle Dubiak.

## Additional Files

Additional file 1: Fig. S1. The heatmap of all differentially expressed genes (DEGs) identified from comparisons between control and Gam1 injected embryos at the three developmental stages (early gastrula, late gastrula, and early neurula, respectively).

Additional file 2: Figure S2. Comparison of microarray data with measurement by qRT-PCR. The fold change between Gam1 and control embryos for the indicated gene as measured by microarray and qRT-PCR are compared with each pair of bars corresponding to early gastrula, late gastrula, and early neurula, respectively. Asterisks indicate samples in which the RNA level was too low to measure by qRT-PCR.

Additional file 3: Figure S3. Volcano plots of differentially expressed genes between Gam1 and control embryos. Fold change (log_2_) is plotted versus statistical significance at (A) early gastrula, (B) late gastrula, and (C) early neurula. Significance values of p < 0.05 (black line) indicates the cut off for accepted differential expression.

Additional file 4: Table S1. Gene Lists Constituting the Biological Processes Most Affected by Loss of SUMOylation Activity.

Additional file 5: Table S2. Biological Processes Associated with Transcription Factors Identified in Network Building.

Additional file 6: Table S3. SUMO Targets Sites and SUMO Interaction Motifs (SIM) in Top Transcription Factors from Network Building.

